# A Novel Proposal for an Index for Regional Cerebral Perfusion Pressure – A Theoretical Approach Using Fluid Dynamics

**DOI:** 10.1101/2021.07.18.452866

**Authors:** Masashi Kameyama, Toshimitsu Momose

**Affiliations:** Department of Diagnostic Radiology, Tokyo Metropolitan Geriatric Hospital and Institute of Gerontology, Tokyo, Japan; Department of Nuclear Medicine, Graduate School of Medicine, The International University of Health and Welfare, Narita, Japan

**Keywords:** positron emission tomography (PET), cerebral blood flow (CBF), cerebral blood volume (CBV), cerebral perfusion pressure (CPP), fluid mechanics

## Abstract

Cerebral blood flow (CBF) / cerebral blood volume (CBV) ratio derived by [^15^O] H_2_O/ CO_2_ and CO positron emission tomography (PET) examination has been used empirically as an index for cerebral perfusion pressure (CPP). However, this equation lacks a theoretical basis and furthermore, has not been confirmed to be proportionate to CPP, as direct measurement of local CPP is not practical.

We have developed a new index for CPP using the Poiseuille equation based on a simple model. Our model suggests that CBF/CBV^2^ is proportionate to CPP and that it is mathematically a more accurate index than CBF/CBV.

## Introduction

Cerebral perfusion pressure (CPP) is the driving force for cerebral blood flow (CBF) and therefore, is an important factor for evaluation of a patient’s cerebral haemodynamic state. However, a noninvasive method for measuring local CPP directly has yet to be developed.

As the CBF/cerebral blood volume (CBV) ratio derived by [^15^O] H_2_O/ CO_2_ and CO positron emission tomography (PET) examination reflected artery patency, CBF/CBV was proposed as an index for haemodynamic reserve [1]. CBF/CBV was further found to be related to oxygen extraction fraction [2] and mean arterial pressure in baboons [3]. Based on these findings, CBF/CBV came to be used as an index for CPP [4, 5]. When CPP decreases, CBV increases and CBF decreases, therefore, CBF/CBV certainly shows some relation to CPP.

However, the index was empirically derived, and has no mathematical basis. Whilst CBF/CBV rises as CPP rises and vice versa, there is no evidence of ratio scale (*i*.*e*. there is no evidence that changes in CBF/CBV is proportional to changes in CPP). In this study, CPP was theoretically derived from CBF and CBV using fluid dynamics.

## Theory

Assume there is one small cerebral region which contains one capillary. The capillary is the sole blood supply for the entire region (Figure 1).

**Figure 1:**
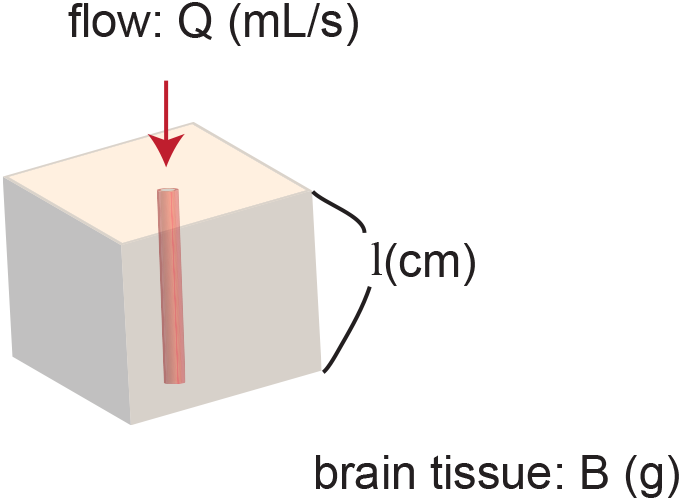
Model of a region of brain. The region contains one capillary. The capillary supplies blood flow to the entire region.

The Poiseuille equation, which can be derived from the Navier-Stokes equations [6] describes incompressible fluid in lamina flow through a long pipe:

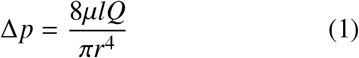

where Δ*p* denotes pressure difference between the two ends, (*i*.*e*. local CPP (Pa)), *µ* is dynamic viscosity (Pa min), *l* is length of capillary (cm), *Q* is volumetric flow rate (mL/min), and *r* is radius of the capillary (cm). CBF and CBV can be calculated as follows:

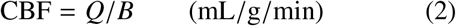

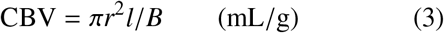

where B denotes local brain tissue weight including the one capillary (g).

Therefore,

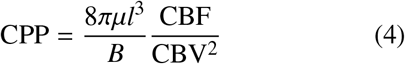

As *µ* is constant and the volume of the brain region perfused by the single capillary is also approximately constant, CPP is proportional to CBF/CBV^2^.

## Method

A simulation was executed to demonstrate how CBF/CBV and CBF/CBV^2^ behave using a standard spreadsheet software, Excel (Microsoft Corporation, Redmond, WA, USA). It was run under the condition that the rate reduction of CBF after the auto-regulation limit (Power’s stage II [7]) was twice the CBV elevation before the limit was reached (stage I).

## Results

CBF/CBV^2^ showed better linearity than CBF/CBV (Figure 2).

**Figure 2:**
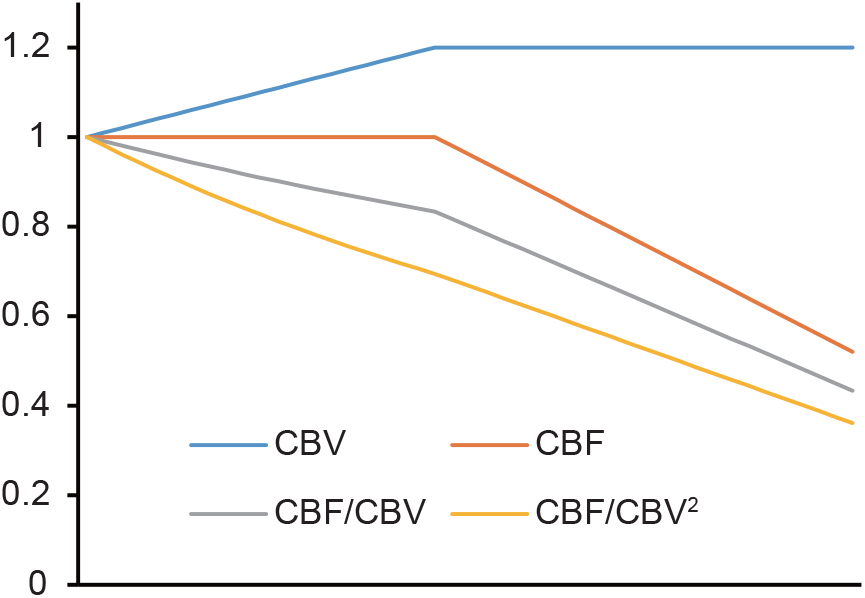
A simulation of CBF/CBV and CBF/CBV^2^.

## Discussion

We have demonstrated theoretically that CBF/CBV^2^ is an appropriate indicator for CPP. CBF/CBV is the reciprocal number of mean transit time [8]. Therefore, it is proportionate to the mean velocity of blood, and certainly relates to CPP. However, our theoretical approach implies that CBF/CBV^2^ would be a better approximation for CPP than CBF/CBV.

The linearity shown in Figure 2 was determined in part by our assumption of CBF reduction being twice that of CBV elavation. However, this is a reasonable assumption considering the fluid dynamics equations above (1, 2, 3, 4). Moreover, a previous study has indicated that lower mean arterial pressure leads to lower CBF/CBV [3].

The Poiseuille equation is applicable under the conditions of laminar flow in a long tube. Thus, our conclusions may not be applicable in situations of turbulent flow. However, the effect of turbulent flow are likely limited as Reynolds number 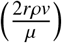 (*v*: velocity of fluid, *ρ*: the density of the fluid) of small capillaries is small.

## Author’s contributions

MK contributed to the conceptualization, creation of theory, and initial draft manuscript preparation. TM advised the project. All the authors discussed the project and have read and approved the final manuscripts.

## Acknowledgements

The authors would like to thank Dr. Natalie Okawa for English language editing of this manuscript.

